# Time-division multiplexing (TDM) sequence removes bias in T2 estimation and relaxation-diffusion measurements

**DOI:** 10.1101/2024.06.03.597138

**Authors:** Qiang Liu, Borjan Gagoski, Imam Ahmed Shaik, Carl-Fredrik Westin, Elisabeth A. Wilde, Walter Schneider, Berkin Bilgic, William Grissom, Jon‐Fredrik Nielsen, Maxim Zaitsev, Yogesh Rathi, Lipeng Ning

## Abstract

**Purpose:** To compare the performance of multi-echo (ME) and time-division multiplexing (TDM) sequences for accelerated relaxation-diffusion MRI (rdMRI) acquisition and to examine their reliability in estimating accurate rdMRI microstructure measures.

**Method:** The ME, TDM, and the reference single-echo (SE) sequences with six echo times (TE) were implemented using Pulseq with single-band (SB-) and multi-band 2 (MB2-) acceleration factors. On a diffusion phantom, the image intensities of the three sequences were compared, and the differences were quantified using the normalized root mean squared error (NRMSE). For the in-vivo brain scan, besides the image intensity comparison and T2-estimates, different methods were used to assess sequence-related effects on microstructure estimation, including the relaxation diffusion imaging moment (REDIM) and the maximum-entropy relaxation diffusion distribution (MaxEnt-RDD).

**Results:** TDM performance was similar to the gold standard SE acquisition, whereas ME showed greater biases (3-4× larger NRMSEs for phantom, 2× for in-vivo). T2 values obtained from TDM closely matched SE, whereas ME sequences underestimated the T2 relaxation time. TDM provided similar diffusion and relaxation parameters as SE using REDIM, whereas SB-ME exhibited a 60% larger bias in the <R_2_> map and on average 3.5× larger bias in the covariance between relaxation-diffusion coefficients.

**Conclusion:** Our analysis demonstrates that TDM provides a more accurate estimation of relaxation-diffusion measurements while accelerating the acquisitions by a factor of 2 to 3.

## Introduction

Diffusion MRI (dMRI)^1^ can probe the microstructure of biological tissues. By leveraging the magnetic gradient, dMRI sensitizes MRI signals to the random microscopic diffusion of water molecules within tissues. MRI relaxometry^2^, is another type of MR contrast mechanism that provides information about the biochemical composition in biological systems by measuring the relaxation processes of water protons. Traditionally, the analysis of dMRI involves data acquisition with a fixed echo time (TE), operating under the assumption that the diffusion process remains independent of T2 (transversal) relaxation. However, the advent of multidimensional dMRI integrates multiple independent experimental parameters into the acquisition strategy, which enables the exploration of correlations between diverse MR contrasts, offering complementary insights into the intricacies of biological tissues by using multi-compartment models.^3,4^ Further, it has been shown that tracing of white matter fiber bundles (such as the superficial white matter) depends on the TE at which the data is acquired.^5,6^

Joint modeling of dMRI with a wide range of TEs, or relaxation-diffusion MRI (rdMRI), has been investigated in several studies^7–12^ to derive novel microstructural measures that enhance our understanding of the underlying biological tissues. These studies include estimating the joint relaxation diffusion distribution (RDD),^7–9^ pre-defining sets of compartments followed by estimating compartmental diffusivities and T2 values,^10^ measuring a compartment-specific T2 value with dMRI data,^12^ estimating fiber-bundle specific T2 values in a tractography framework^5,6^ and estimating the joint moments of relaxation and diffusion.^11^ However, a notable challenge hindering the use of rdMRI techniques is the prolonged scan time required for data acquisition. Typically, this involves repeating a diffusion scan with varying TEs while keeping the repetition time (TR) and the diffusion time constant among the repetitions for T2-diffusion studies to mitigate the influence of T1 (longitudinal) relaxation effects. For instance, acquiring 6-TE dMRI data using single-band single-echo (SE) EPI (which takes 12-min considering 3 *b*-values, 25 diffusion directions each, and whole brain coverage) can take 1 hour 12 min. Consequently, the rdMRI acquisition process becomes inefficient due to the idle time in the sequence.

One efficient scheme to accelerate the rdMRI sampling is using the multi-echo (ME) spin-echo sequence.^3,13-14^ By applying multiple 180-degree refocusing radiofrequency (RF) pulses, the ME sequence obtains multiple echoes at different TEs that share the same diffusion preparation in one TR. Unlike the conventional SE sequence, the ME sequence maximizes TR utilization by incorporating additional echoes, minimizing idle time. Despite its potential to enhance rdMRI acquisition efficiency, concerns persist regarding image signal biases in the following echoes of ME attributable to transmit field (B1+) inhomogeneity and coarse slice profiles stemming from imperfect refocusing pulses,^15^ as documented in prior studies employing the ME-SE sequence for accelerating T2 mapping^16^. Yet, the accuracy of using ME to expedite rdMRI has not been investigated and evaluated.

The time-division multiplexing (TDM)^17,18^ sequence is an alternative strategy to accelerate rdMRI scans by acquiring multiple slices at different TEs while each slice maintains the SE pattern, avoiding multiple refocusing pulses per slice. Within a given TR, TDM sequence effectively interleaves and rearranges sequence event blocks—encompassing RF excitation, refocusing, and readout for up to three slices. Echo-time shifting gradients are employed to mitigate interference between slices and coupling with diffusion gradients. Notably, through integration with the simultaneous multi-slice (SMS)^19^ technique, our previous research has demonstrated that TDM3 (acceleration factor of 3) with an SMS multi-band (MB) factor of 2 can achieve a total acceleration factor of up to 6× compared to the conventional single-band (SB-) SE sequence, all while maintaining image quality. Building upon this foundation, our recent study^18^ has employed MB2-TDM3 to acquire multi-TE dMRI data, wherein relaxation-regressed dMRI data were estimated. This marks a crucial step towards leveraging TDM to accelerate and benefit the study of rdMRI. However, a direct comparison of these sequences (TDM vs ME) on their ability to accurately estimate T2 and rdMRI measures has not been done.

In the current study, we systematically compared the performance of employing TDM and ME methods to accelerate rdMRI acquisition and examined the reliability of using TDM and ME sequences to derive rdMRI microstructure measures. We implemented the ME, TDM, and reference SE sequences (incorporating various TEs) in Pulseq^20-23^ (an open-source and vendor-neutral pulse sequence development platform), in both SB- and MB2-formats, to accurately match sequence parameters. To further investigate the impact of slice profile on ME sequences, in addition to the standard Sinc RF pulses, we also explored alternative versions, implementing Shinnar–Le Roux (SLR) pulses^24^ with different time-bandwidth products (TBWPs) for the SB-ME and SB-SE sequences. Our study used a phantom with various T2 values and a healthy human subject, with comparisons using several metrics and measures:

### Phantom Analysis

We compared the image intensities of the three sequences (in both SB- and MB2-formats) and reported differences using the normalized root mean squared error (NRMSE). Furthermore, we calculated and compared the estimated T2 values between the six sequences.

### Slice Profile Improvement

We compared the image intensities from SB-ME against SB-SE, both with SLR RF pulses, to investigate whether enhancing the slice profile would mitigate signal biases in the ME sequence.

### In-vivo Brain Scan

We examined image intensities and quantified differences between ME, TDM, and SE sequences. Furthermore, we applied methods developed in our previous work to assess sequence-related effects on microstructure estimation. Specifically, we utilized the relaxation diffusion imaging moment (REDIM)^11^ method to estimate the joint moments of T2 relaxation and diffusion coefficients. Additionally, on the MB2 data, we employed the maximum-entropy relaxation diffusion distribution (MaxEnt-RDD)^8^ method to estimate the joint distribution of relaxation and diffusion coefficients in each voxel in the hippocampus.

All the sequences performed in this work and the corresponding reconstruction code are accessible at: https://github.com/QiangLiu0310/Pulseq_TDM_acc_rdMRI.

## Methods

### Sequence implementation and scan protocol

This study was conducted following approval from the local Institutional Review Boards (IRBs). All experiments were performed on a clinical 3T scanner (Prisma, software version XA30, Siemens Healthineers, Erlangen, Germany). A phantom with various T2 values and a healthy male subject were scanned.

We implemented SE (reference), ME, and TDM sequences with no slice acceleration (SB) and multi-band acceleration (MB=2). A schematic illustration of these sequences is provided in Figure 1(A). For the ME sequence,^14,25^ after the first refocusing RF pulse and the corresponding EPI readout, a second refocusing RF pulse was applied to the same slice, and the second echo was acquired. The third echo was generated and collected following the same pattern. The TDM sequence,^17,18^ on the other hand, employed the echo time-shifting gradients, collecting three echoes at different TEs from three different slices. Since the MB2-TDM3 sequence provided a higher total acceleration factor of 6 (MB2 × TDM3), it had the shortest minimum TR among the three sequences. To remove TR-dependent effects and make the total scan time feasible for a human scan, we implemented a special slice loop for the SE and ME sequences to match their TRs with the TDM sequence, as shown in Figure 1(B). Each loop of the SE and ME sequences only included a subset of the total slices so that they could have the same TR as the TDM sequence. The sequences were repeated with a shifted set of slices to obtain whole-brain scans. These sequences were programmed and developed using the vendor-neutral platform Pulseq,^20^ to ensure identical readout gradients.

**Figure 1.**
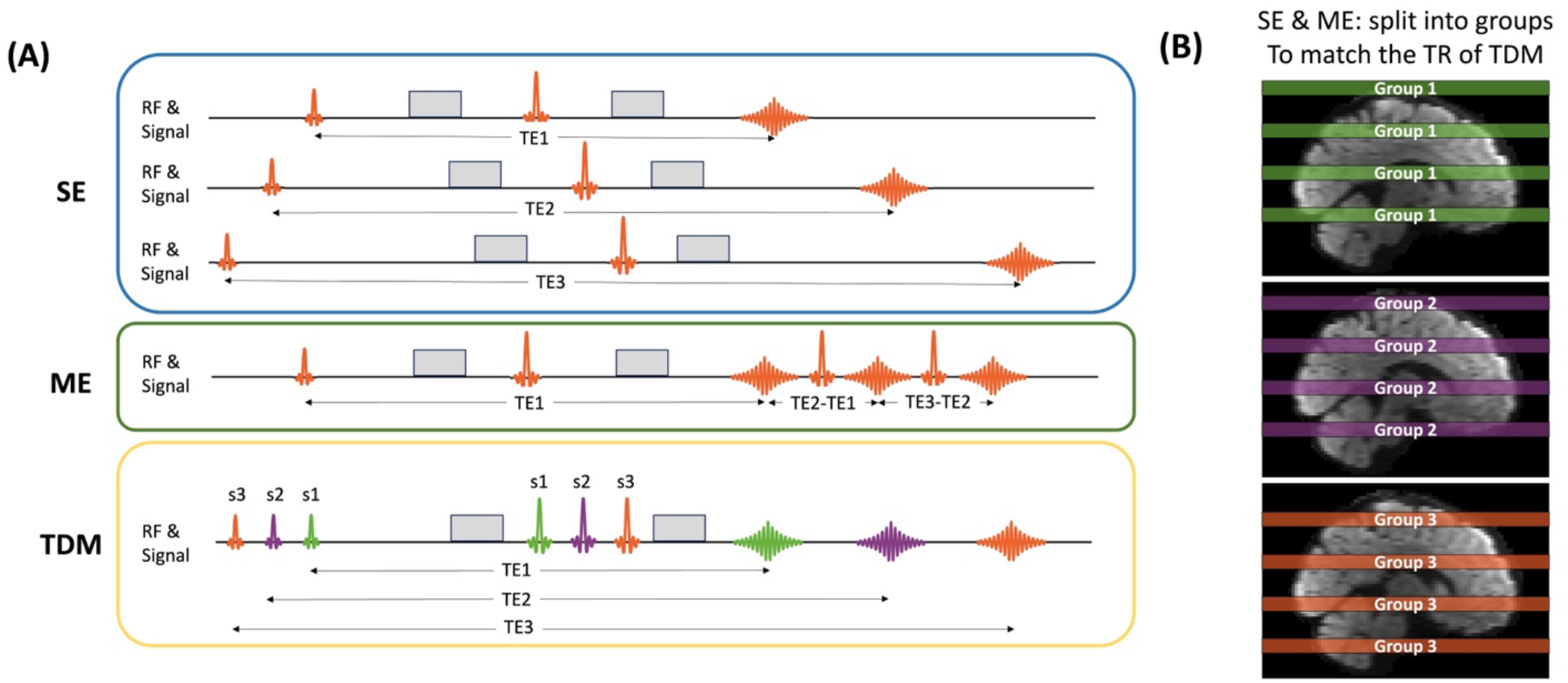
(A) Schematic diagrams illustrating the single-echo (SE-) EPI, multi-echo (ME-) EPI, and time-division multiplexing (TDM) sequences. In the ME sequence, three echoes were acquired at the same slice, while in TDM, three echoes at different slices were acquired at different echo times (TEs). The diffusion times were kept consistent across all sequences to mitigate diffusion time-dependent effects. (B) For SE and ME sequences, a specialized acquisition scheme was implemented to synchronize the repetition time (TR) with TDM. The total slices were partitioned into groups (two for single-band (SB) and three for multi-band 2 (MB2), based on the total slice number), with each loop encompassing only a subset of the slices.

The phantom and the in-vivo scans shared the same scan parameters: the SE, ME, and TDM sequences were performed with TR=2500 ms, TEs=[70,82,95,107,120,132] ms, echo spacing=0.48 ms, FOV=220 × 220 mm^2^, in-plane resolution=2.5 × 2.5 mm^2^, partial Fourier factor=6/8, in-plane GRAPPA^26^ factor=3, slice thickness=2.5 mm. The duration of the excitation Sinc RF pulse was 3 ms, with a TBWP of 4, while the 180-degree refocusing Sinc RF pulse had a duration of 5 ms, TBWP=4. The ME sequence was performed in the format of three-echo, while the TDM sequence was with three-slice at 3 different TEs. To acquire rdMRI data with 6 TEs using ME and TDM sequences, two ME and two TDM sequences were performed with two TE groups: TE_group1_=[70,95,120] ms and TE_group2_=[82,107,132] ms. In the in-vivo scans, aside from the *b*=0 acquisition, *b*-values of [1000,2000,3000] s/mm^2^ were obtained for all three sequences for TE_group1_, and *b*-values of [500,1500,2500] s/mm^2^ for TE_group2_. In total, rdMRI data of 6 TEs and 6 b-values were sampled in a complementary diffusion-relaxation 2D sampling space. For all *b*-values and TEs, 25 diffusion directions were acquired. To avoid diffusion time-dependent effects for SE, ME, and TDM, we set *δ*=11.9 ms and Δ=47.1 ms for all TEs. A separate *b*=0 volume with opposite phase-encoding gradients was performed for all three sequences at each TE. A field-of-view (FOV) matched fully sampled Pulseq 3-shot EPI was acquired as the reference scan data before each sequence. T1-weighted structural images were acquired for the subject as the anatomical reference using a 3D T1-weighted image magnetization-prepared rapid gradient echo^27^ (MPRAGE) sequence.

A separate single-band experiment utilizing different SLR RF pulses was conducted on the phantom. Three sequences were implemented with the above scan parameters: SB-SE sequence with SLR pulses and TBWP of 4, and SE-ME sequence with SLR pulses with TBWP values of 4 and 6, respectively. RF power calculations for the refocusing RF pulses in these sequences were performed using the Pulseq function *calcRfPower*, and three parameters were subsequently reported: total energy, peak power, and RMS (root-mean-square) B1 amplitude.

For the single-band experiment, 18 slices were acquired in an interleaved pattern. We used a partial field of view, keeping in mind the long scan time (2 hours) required to obtain all scans (SE, TDM, ME) during the same session. To align the TR with the TDM sequence, these 18 slices were divided into two subsets for SE and ME sequences (as depicted in Figure 1(B)). The experiments were repeated with MB sequences (MB=2) to examine if the MB RF pulses and the blipped-caipi^19^ readout lead to additional differences between the sequences. In the MB2 experiments, 60 slices were collected in an interleaved manner, similar to the schematic representation in Figure 1(B), where three subgroups were created for SE and ME sequences.

### Image reconstruction and post-processing

The k-space raw data were initially converted to ISMRM-RD format^28^ for all sequences, and custom MATLAB scripts were used to reconstruct the images. GRAPPA^26^ was utilized to reconstruct the SB data, whereas split slice-GRAPPA^29^ was employed for MB2 data, and GRAPPA for the in-plane GRAPPA factor=3 image reconstruction. Subsequently, FSL^30^ *TOPUP*^31^ and *eddy*^32^ were used to correct the susceptibility-induced distortion and eddy current effects at each TE separately. To bring all the acquired images from multiple sessions into a common space, rigid registration was performed using Advanced Normalization Tools^33^ (ANTs) for all the dMRI images at each TE to the images from SB/MB2-SE TE=70 ms, for both phantom and in-vivo scans. *FreeSurfer*^34^ was used to parcellate the brain using the Deskian-Killiany atlas on the T1-MPRAGE images, and the outputs of *FreeSurfer* were registered to the diffusion space by ANTs.

### Data analysis on the phantom data

For the phantom data (both SB and MB sequences), the NRMSE in signal intensity of *b*=0 images between TDM, ME, and standard SE sequences (considered as gold standard) was computed and compared to examine the relative changes in image intensity. Furthermore, the T2 value was estimated from the 6 TE images using a mono-exponential fitting model employing the *fminsearch* function in the MATLAB optimization toolbox. Six regions of interest (ROIs) of different T2 values, each with an area of 21 pixels, were manually drawn^34^ within one slice on the phantom using *3D Slicer*^35^.

### Data analysis on the in-vivo data

For the in-vivo data, first, the NRMSE was calculated in the rdMRI data between TDM, ME, and the SE sequences (with Sinc RF pulses of TBWP=4) for both SB and MB variants. Next, we examined sequence-dependent effects on image intensity and rdMRI microstructural measures. By comparing results with different combinations of SB, MB, TDM, and ME techniques, we can identify the most reliable method for rdMRI studies. First, the NRMSE of TDM and ME sequences compared to SE was calculated for both MB and SB data. Different from the phantom experiments, the NRMSE was computed using direction-averaged signals at all b-values. The direction average method can reduce the impact of measurement noise for a more accurate analysis.

Moreover, we applied the methods developed in our previous works^8,11^ to compute rdMRI measures to examine sequence-related effects on microstructure estimation. First, we used the relaxation diffusion imaging moment^11^ (REDIM) method to estimate the joint moments of T2 relaxation and diffusion coefficients. The coefficients were estimated by solving the least-square fitting problem with linear constraints using the *lsqlin* function in Matlab using direction-averaged signals. The method provides the mean relaxation rate <R_2_>, mean diffusion coefficients <D>, and the covariance between the relaxation and diffusion coefficients CRD. Moreover, we applied the maximum-entropy relaxation diffusion distribution methods^8^ (MaxEnt-RDD) to estimate the joint distribution of relaxation and diffusion coefficients in each voxel. The MaxEnt-RDD method provides an approach to identifying the microstructural properties without assuming a specific number of tissue components. Our previous work^35^ has shown that MaxEnt-RDD is sensitive to detecting brain abnormalities related to temporal lobe epilepsy that cannot be detected using structural MRI and relaxation MRI. Differences in MaxEnt-RDD functions between the right and left hippocampus provide information to lateralize epileptic lesions. To this end, we computed the average MaxEnt-RDD from the three sequences in the left and right hippocampus regions. We note that other multi-compartment models^10,36^ are available for microstructure analysis using rdMRI. Different from these methods, the chosen REDIM and MaxEnt-RDD methods can be solved by using convex optimization algorithms that can avoid issues related to local minima to ensure reliable comparison of microstructure measures between different sequences.

## Results

### Phantom Results

Figure 2 depicts the signal intensity differences between *b*=0 images obtained from SE, ME, and TDM sequences (with Sinc RF pulses of TBWP=4). Figure 2(A) shows the images from the SB sequences, while Figure 2(B) displays the MB images. SB-ME and MB-ME resulted in different signal intensities of TE=95 ms and TE=120 ms from the SE sequences, whereas SB-TDM and MB-TDM show similar signal levels compared to SE sequences at each TE. The signal biases from the ME sequences are further depicted in the NRMSE maps: the SB-TDM had 3.6× and 4.6× lower NRMSEs for 2^nd^ and 3^rd^ echoes than the SB-ME variant (as shown in the mean NRMSE values in Figure 2(A)); besides TE=70 ms, slightly higher biases could be found for MB-ME sequence at TE=95 ms and TE=120 ms compared to SB variant, where 2.9× and 3.2× lower NRMSEs were achieved by MB-TDM for the 2^nd^ and 3^rd^ echoes, respectively.

**Figure 2.**
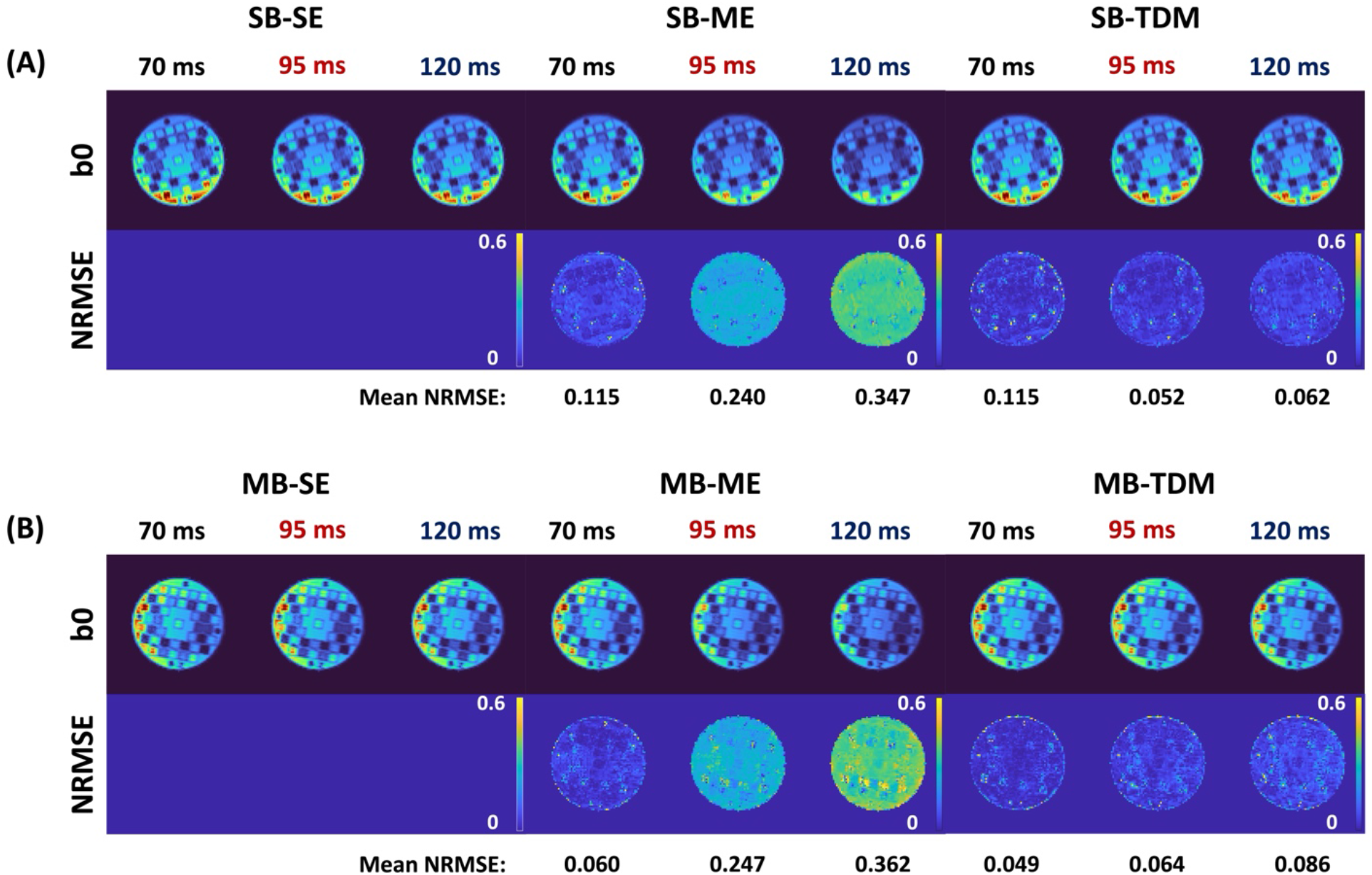
Phantom experiment: Signal intensity differences of the *b*=0 images on the phantom at three different echo times (TEs) from SE, ME, and TDM sequences (with Sinc RF pulses with time-bandwidth product (TBWP)=4). Figure 2(A) presents images from single-band (SB) sequences, while Figure 2(B) displays multi-band (MB) images. The normalized root mean squared error (NRMSE) was computed between the ME, TDM, and SE images, with the mean NRMSEs within the phantom depicted.

Figure 3(A) displays signal intensity and NRMSE maps acquired from *b*=0 images using SB-SE sequence employing SLR pulses with TBWP=4, and SB-ME sequences employing SLR pulses with TBWP values of 4 and 6, respectively. When compared to SB-ME with Sinc pulses, reductions in signal intensity biases were observed with the SLR pulses, as illustrated by the NRMSE maps. Specifically, for SB-ME SLR with TBWP=4, the mean NRMSEs of TE [70, 95, 120] ms were [0.034, 0.063, 0.127] respectively, while for the TBWP=6 pulses, NRMSEs were further diminished and comparable to our SE-TDM, yielding [0.090, 0.056, 0.062]. Figure 3(B) illustrates the RF power of these three refocusing RF pulses and their ratios against the Sinc refocusing pulse with TBWP=4. The total energy and peak power of the SLR pulse were 2.34× and 3.5× compared to those of the Sinc pulse, respectively. As anticipated, increasing the TBWP resulted in a further increase in RF power, causing a 3.63× and 8.46× increase in total energy and peak power required, respectively.

**Figure 3.**
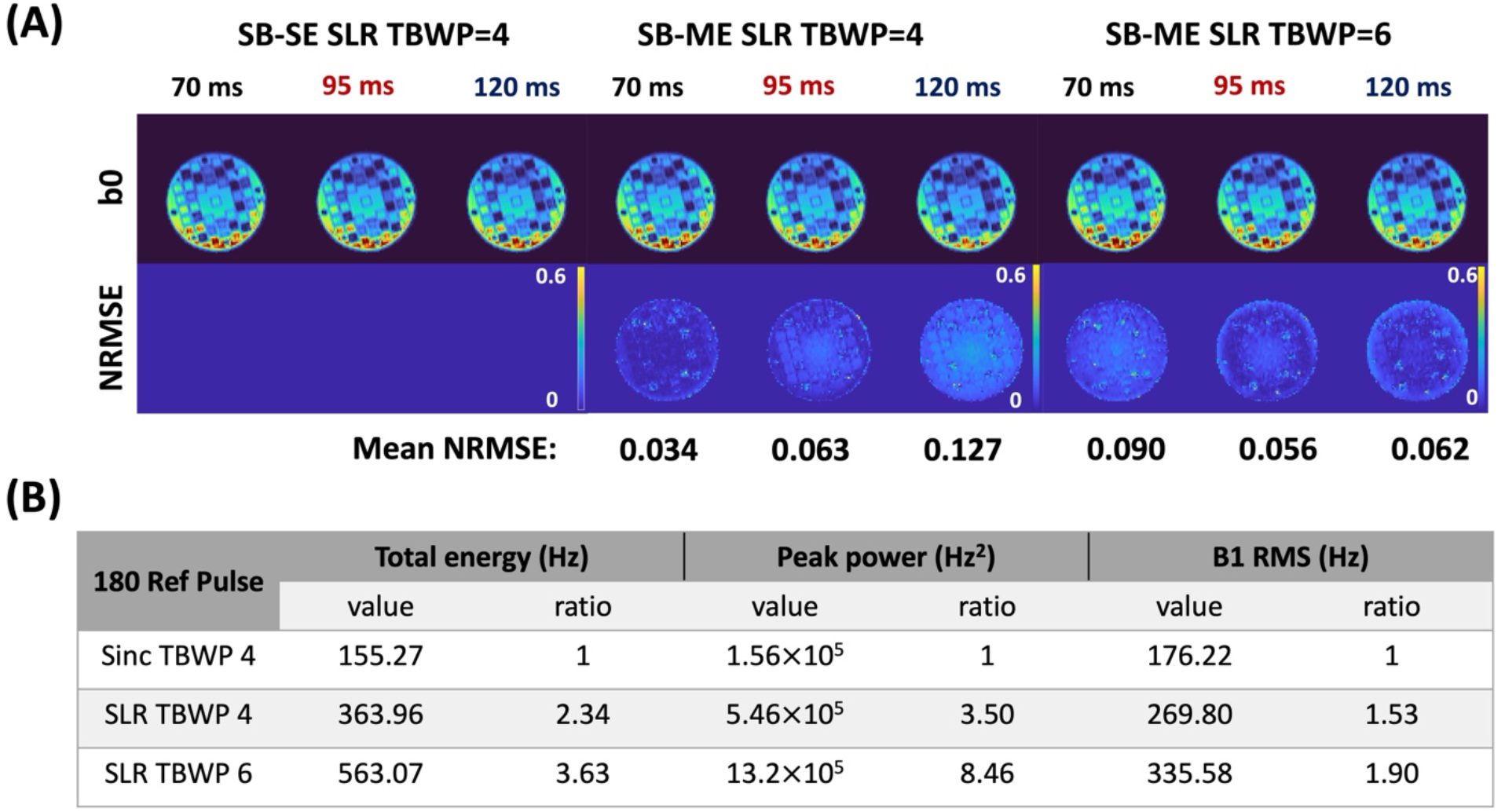
Phantom experiment: (A) The signal intensity of the b=0 images acquired with SB-SE sequence using SLR pulses with a time-bandwidth product (TBWP)=4, SB-ME employing SLR pulses with TBWP=4 and 6. The NRMSEs between ME and SE sequences are shown in the second row, and the mean NRMSEs within the phantom are displayed below. (B) The RF power of the 180-degree refocusing pulses. Three RF pulses are evaluated, which are Sinc pulse with a TBWP=4, SLR pulse with TBWP=4, and 6. The RF power calculation considers total energy, peak power, and B1 root-mean-square (RMS). The RF power values and their ratios compared to the Sinc pulse with TBWP=4 are presented.

Figure 4 shows the T2 estimation of the phantom. Figure 4(A) shows the T2 map estimated from the 6 echoes using the SB-SE sequence, with six ROIs drawn using 3D *Slicer* displayed. Figure 4(B) shows the T2 values estimated from SB and MB variants of SE, ME, and TDM sequences. Notably, T2 values obtained from TDM closely matched those from SE variants for both SB and MB cases. However, the ME methods yielded underestimated and biased T2 estimations.

**Figure 4.**
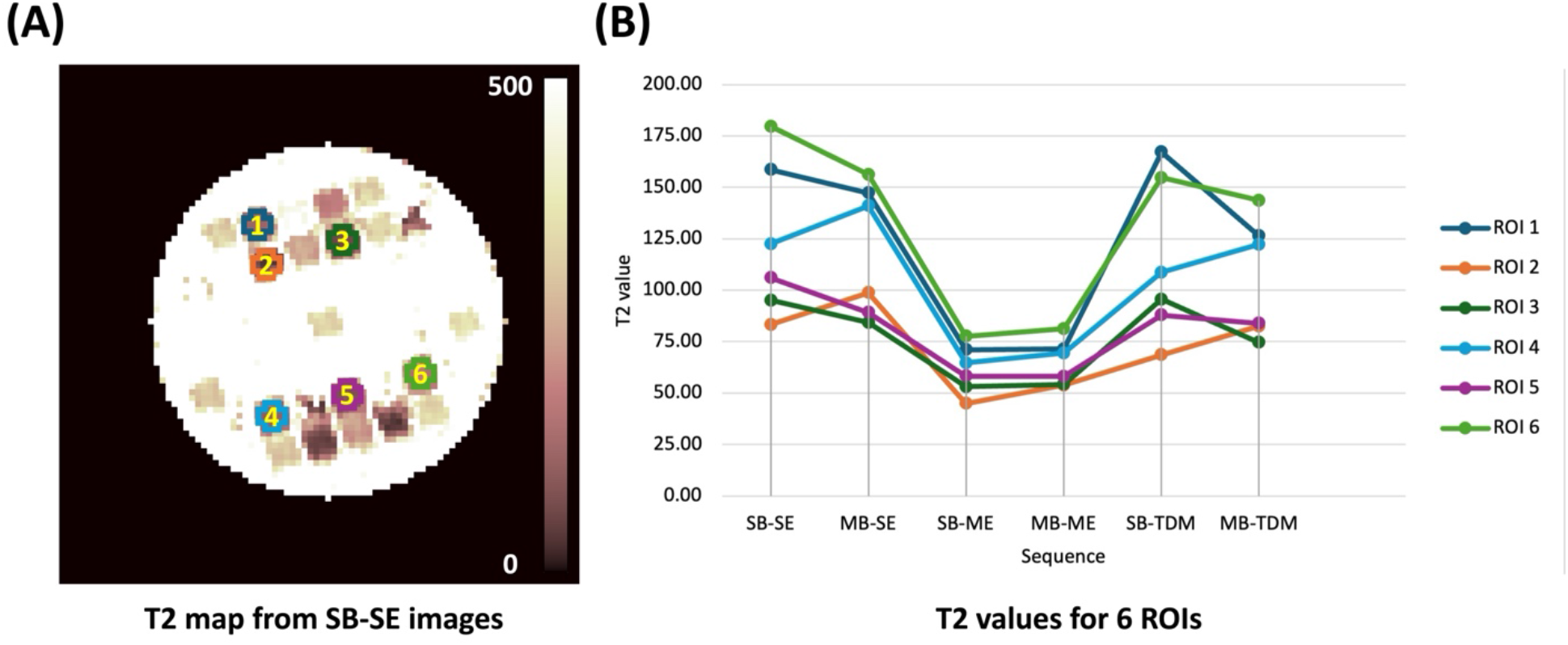
Phantom experiment: (A) The T2 map estimated from 6-TE images acquired using SB-SE sequence with Sinc pulse with TBWP=4. Six region-of-interests (ROIs) were manually drawn using *3D Slicer* and shown. (B) The T2 values were estimated from 6 different TEs using SE, ME, and TDM sequences in both SB and MB formats. Six ROIs with variable T2 values were included.

### Results on in vivo data

Figure 5 displays a representative slice of the mean DWI of the in-vivo scan. Consistent with the phantom experiment, the mean DWIs of the 2^nd^ and 3^rd^ echoes acquired from ME sequences exhibited lower signal levels compared to standard SE variants. Furthermore, the biased estimates are reflected in the calculated NRMSE maps. Employing TDM reduced the NRMSE by half compared to ME variants, with SE variants serving as references, as shown in the last line in Figure 5 (A) and (B).

**Figure 5.**
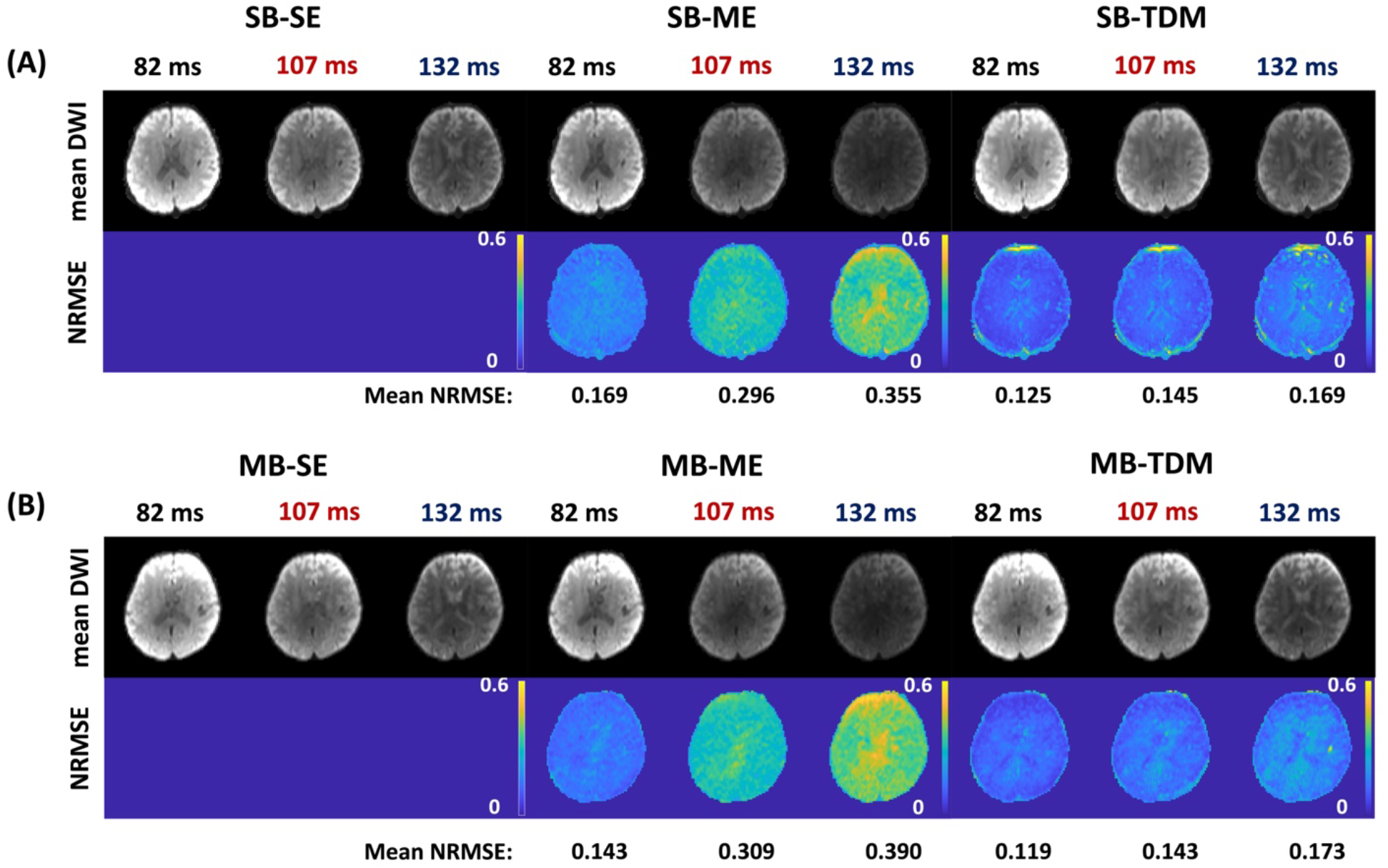
In-vivo experiment: Signal intensity differences of the direction-averaged (mean DWI) images with *b*-value=500 s/mm^2^ at three different echo times (TEs) from SE, ME, and TDM sequences (with Sinc RF pulses of a time-bandwidth product (TBWP)=4). Figure 5(A) presents images from SB sequences, while Figure 5(B) displays MB images. The NRMSE was computed between the ME, TDM, and SE images using the direction-averaged images at all the *b*-values, with the mean NRMSEs within the brain ROI depicted below.

Figure 6 presents <D> (Figure 6(A)), <R_2_> (Figure 6(B)), and CRD maps (Figure 6(C)) derived from the REDIM method using the SB rdMRI data, along with the differences observed between each sequence and the SB-SE sequence. The quantitative metrics for all the measures provided by SB-TDM closely align with those of the reference SB-SE sequence, as can be seen in the difference maps. Although the average diffusivity map calculated from the SB-ME sequence deviated slightly from the gold standard SB-SE method, the <R_2_> map derived from SB-ME exhibited a 60% larger bias compared to the reference SB-SE scan. Additionally, the CRD map from SB-ME exhibited a bias yielding higher positive values in the white matter region and in cerebrospinal fluid (CSF) compared to the reference.

**Figure 6.**
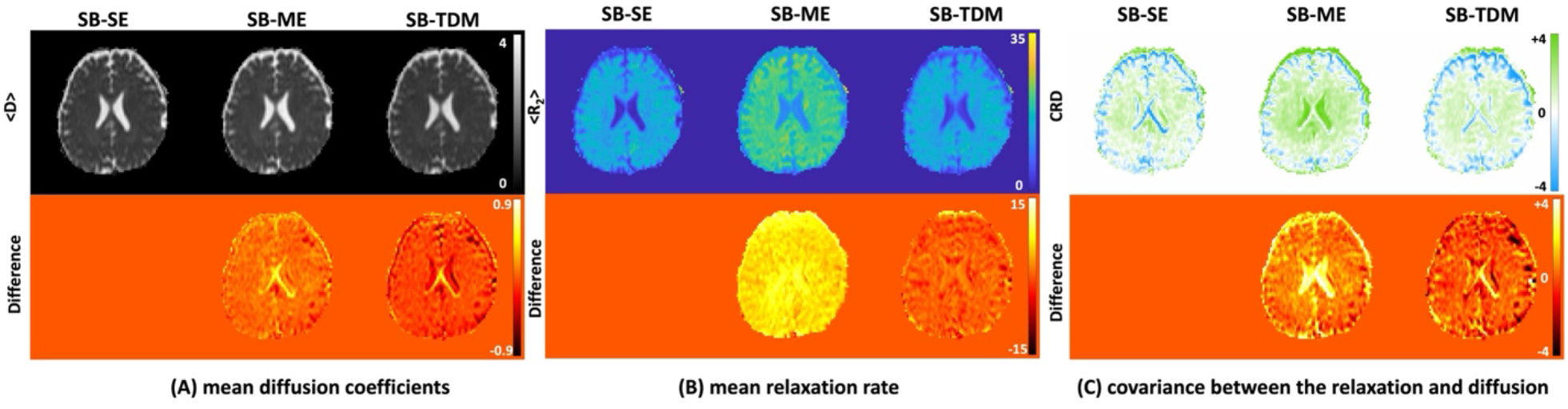
The joint moments of T2-relaxation and diffusion coefficients estimated using the relaxation diffusion imaging moment (REDIM) method with the rdMRI data acquired by SB-SE, SB-ME, and SB-TDM sequences. (A) The mean diffusion coefficients <D>, (B) mean relaxation rate <R_2_>, and (C) the covariance between the relaxation and diffusion coefficients CRD. The top row displays the measurements derived from the REDIM method, and the bottom row shows the differences between the measurements from SB-ME, SB-TDM, and SB-SE data.

Figure 7(A) shows the averaged <R_2_> values in three brain regions (subcortical gray, cortical gray, and white matter) using MB-SE, MB-ME, and MB-TDM sequences. MB-TDM retained similar mean relaxation rates as the reference MB-SE measurements, whereas MB-ME showed higher <R_2_> in all three regions. Figure 7(B) provides the averaged CRD values in some regions of the deep brain areas for three sequences. For MB-TDM, the average CRD values in these regions were similar to MB-SE, while MB-ME demonstrated biases and presented higher values. The mean absolute difference between CRD values for MB-TDM vs MB-SE across 12 brain regions was 3.5× lower than that of MB-ME vs MB-SE (0.54 and 1.90, respectively). Moreover, the Pearson correlation coefficient between MB-TDM and MB-SE was 0.9629 (*p*<0.01), while the correlation between MB-ME and MB-SE was 0.159 (*p*=0.6216).

**Figure 7.**
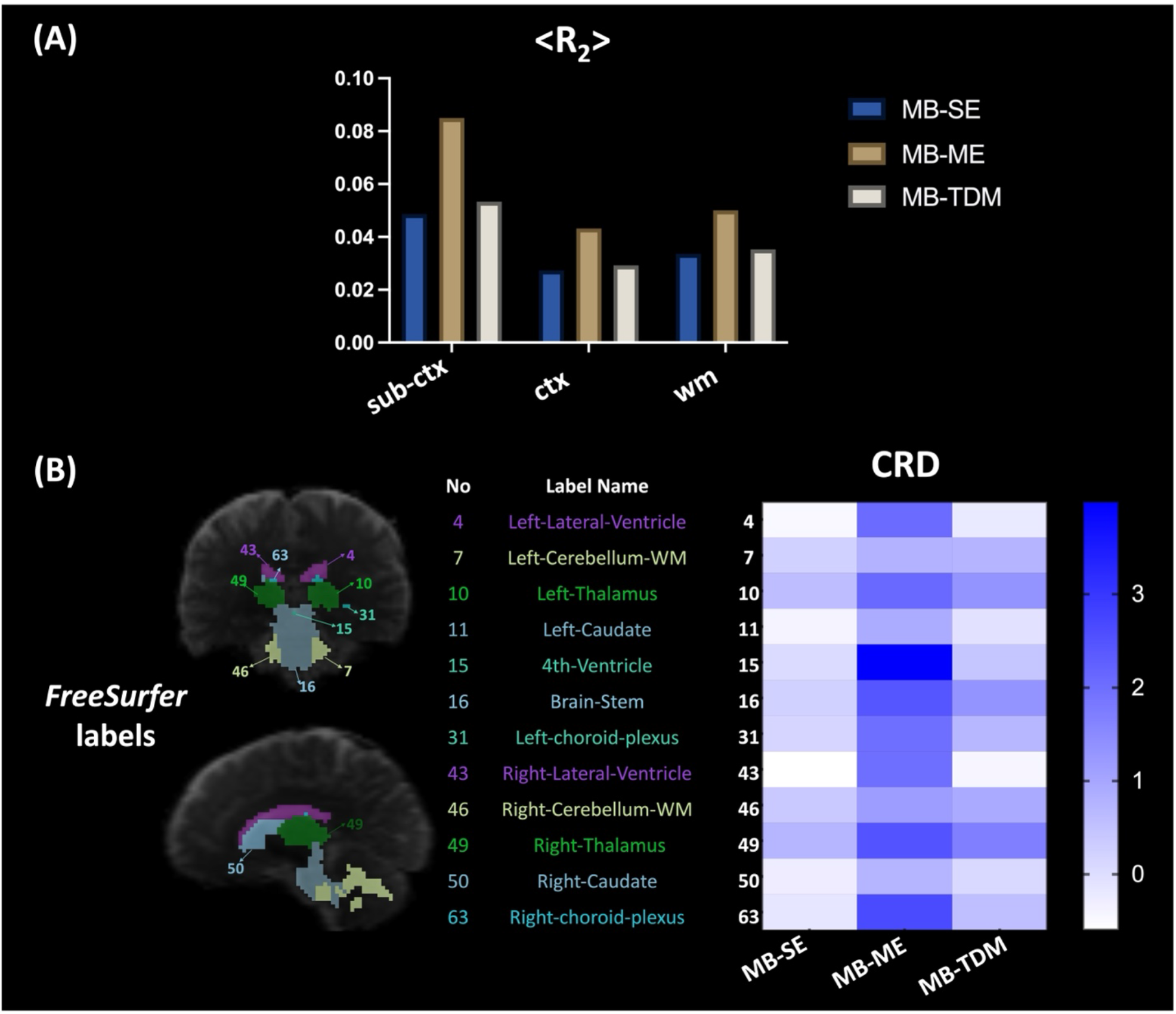
Diffusion-relaxometry parameters were computed using REDIM on the MB rdMRI data. (A) The average <R_2_> values in subcortical gray (sub-ctx), cortical gray (ctx), and white matter (wm) regions using the rdMRI data acquired by MB-SE, MB-ME, and MB-TDM sequences. (B) *FreeSurfer* labels in some deep brain areas are depicted on the *b*=0 image (left column), with the brain regions listed in the middle column, the average CRD values calculated from the three MB sequences displayed on the right.

Figure 8(A) illustrates the parcellation of the hippocampus by *FreeSurfer*. Figure 8(B) shows the RDD function in one voxel from the rdMRI data of the MB-SE sequence, where three components were detected. Figure 8 (C-E) shows the averaged RDD function from all the voxels in the hippocampus using the rdMRI data acquired from MB-SE, MB-ME, and MB-TDM, respectively. The RDD of MB-ME data exhibited a large bias, especially in the area containing low diffusivity and R_2_ values, as shown in the zoom-in figures. MB-TDM, however, retained a distribution similar to MB-SE.

**Figure 8.**
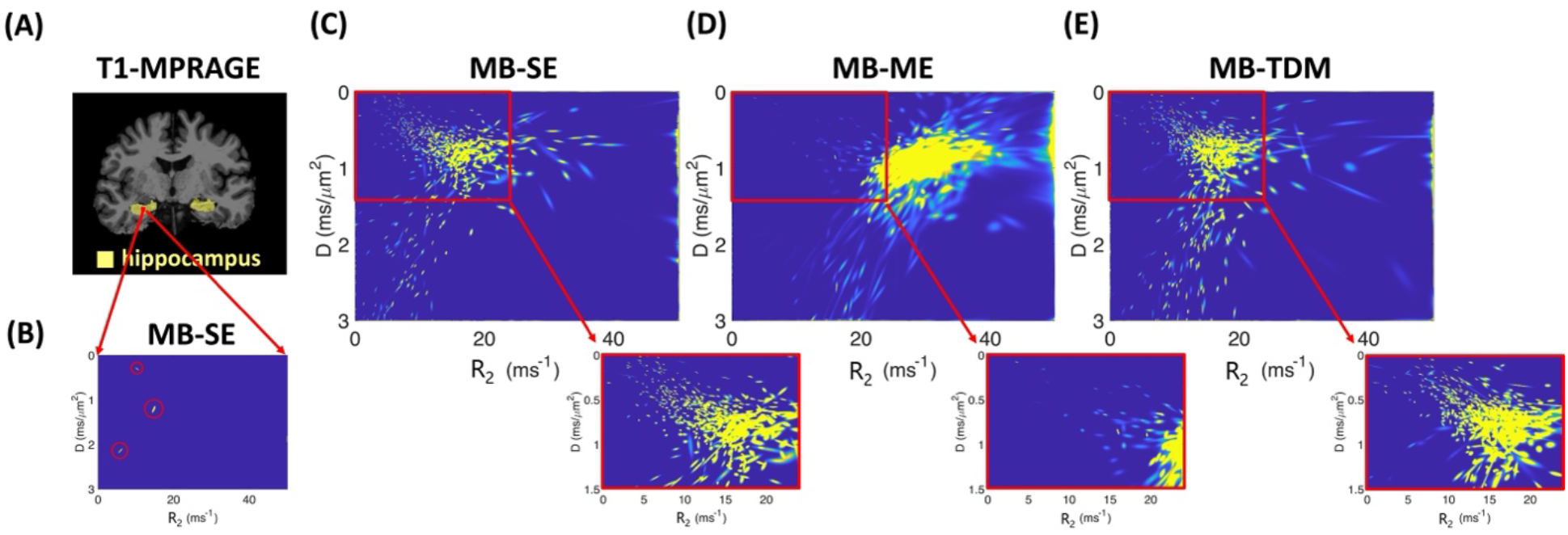
The voxel-wise joint distribution of relaxation and diffusion coefficients estimated by maximum-entropy relaxation diffusion distribution method (MaxEnt-RDD) in the hippocampus. (A) The parcellation of the hippocampus by *FreeSurfer* on the T1-MPRAGE image. (B) The RDD function in one voxel was derived from the rdMRI data using the MB-SE sequence, where three components were detected. (C-E) shows the average RDD function from all the voxels in the hippocampus using the rdMRI data acquired with MB-SE, MB-ME, and MB-TDM sequences, respectively.

## Discussion

In this study, leveraging the flexibility afforded by the Pulseq sequence development platform, we implemented two advanced dMRI sequences, TDM and ME, and systematically assessed their feasibility to accelerate rdMRI acquisitions. Despite both TDM and ME offering the potential to reduce scan time by up to 2 to 3× compared to the standard SE acquisition, our findings indicate that images acquired with the ME sequence exhibit significantly larger intensity biases than those acquired with TDM. Specifically, the intensity biases in the ME sequence were approximately four times higher on the phantom and twice as high in vivo, as indicated by the NRMSEs. Consequently, the ME sequence dramatically underestimated the T2 decay rates across the brain. We further examined the sequence-related effects on microstructure estimation in in-vivo data, and results using the REDIM and MaxEnt-RDD methods proved that TDM provided more reliable estimates for all rdMRI measures.

The image intensity biases observed in the 2^nd^ and 3^rd^ echoes from the ME sequence can be attributed to poor slice profiles. Ideally, after the first TE, the transverse magnetization within a certain slice begins to dephase, and subsequently applied refocusing pulses rephase the bulk magnetization and generate subsequent echoes. However, in practice, due to the non-ideal slice-selection performance of the refocusing pulses, the slice boundaries are not fully refocused, resulting in a drop in signal level. This effect accumulates over subsequent TEs, as evidenced by the increasing NRMSEs. Our study demonstrates that improving the slice profile can alleviate this signal bias. Specifically, on the phantom, we achieved similar NRMSEs with the SB-ME sequence using SLR RF pulses with TBWP=6, comparable to those of SB-TDM with Sinc pulses. However, due to the high RF power induced by the SLR pulse (approximately 8.5× the peak power of the Sinc pulse), direct application in in-vivo rdMRI scans, particularly with SMS combined, may not be practical and create discomfort to patients during long scans. One way to reduce the requirement for increased power could be to use advanced techniques such as root-flipped^37^ or Variable-Rate Selective Excitation^38,39^ (VERSE) to reduce the peak power of the SLR pulses. However, using VERSE would still not create the ideal refocusing slice profile as in the SB case, especially when B1+ and B0 inhomogeneity is aggravated.^40^ Another potential solution is exciting a thicker slice when applying the refocusing RF pulse than the excitation RF pulse^41,42^, but this could introduce spin-history effects and thus prolong the TR. In contrast, TDM acquires multiple slices at different TEs to avoid multiple refocusing pulses per slice, achieving intensity levels close to SE acquisitions. We also note that in Figure 5(A), a region with high NRMSE values can be observed in the frontal area in all three TEs of SE-TDM, which might be related to the misregistration due to the motion during the long scan.

The T2 estimation in the phantom in our study highlights that compared to ME sequences, TDM demonstrates superior reliability in T2 estimation, as shown in the T2-value plot. Furthermore, we observed differences in T2 values measured using the SB-SE and the MB-SE variants, potentially attributable to differing signal-to-noise ratio (SNR) levels influenced by the g-factor. This finding suggests that employing SB-TDM as a slice-acceleration method for rdMRI data acquisition may lead to more reliable T2 estimation.

The joint relaxation and diffusion distribution (RDD) estimation by our MaxEnt-RDD method for the MB-ME data was biased and missed some components in the region of lower R_2_ and lower diffusivity values, while the distribution from MB-TDM was consistent with that of MB-SE. This finding suggests that the ME sequence might not be accurate in separating multiple compartments. In contrast, TDM maintains the sensitivity of detecting heterogeneous tissue microstructure, which could potentially benefit the detection of some diseases. For example, our recent work showed the feasibility of lesion detection for a patient with sMRI-negative epilepsy leveraging the MaxEnt-RDD method with the rdMRI data from MB-TDM.^36^

It is pertinent to mention that beyond the ME and TDM sequences explored in our study, several advanced sequences hold promise for accelerating rdMRI or similar multi-dimensional MRI acquisitions. One such method involves combining multidimensional MR fingerprinting (mdMRF) with b-tensor encoding^43^, enabling the simultaneous quantification of relaxation and diffusion from a single scan. In contrast to diffusion-weighted acquisition as employed in our study, mdMRF utilizes diffusion-prepared SSFP. However, the diffusion-preparation module is more sensitive to phase errors induced by eddy currents or motion compared to standard diffusion-weighted acquisition, leading to signal voids and shading artifacts.^44, 45^ Additionally, an efficient T1, T2, and ADC mapping technique has been developed using MR Multitasking.^45^ Nonetheless, the reconstruction time of this method is prolonged due to the imaging model containing three spatial dimensions and four dimensions indexing the T1/T2/b-value/diffusion-direction variables. Other notable acquisition techniques include ACE-EPTI^46^, which offers distortion-free, blurring-free, and time-resolved ME images, and ZEBRA^47^, enabling simultaneous sampling of the three-dimensional (T1-T2*-diffusion) acquisition parameter space. These diverse approaches showcase the ongoing efforts to enhance multi-dimensional MRI acquisition, each presenting unique advantages and limitations.

This study also has some limitations worth noting. The spatial resolution of the rdMRI is 2.5 mm (isotropic voxel size), which might impact the performance of the ME-RDD method as it is sensitive to noise and is limited by the minimum TEs we can achieve on a clinical scanner. Our previous study has shown that on MAGNUS, equipped with a high-performance gradient system, isotropic 1.5 mm rdMRI with a minimum TE of 45 ms can be obtained with our TDM sequence.^48^

It is noteworthy that the minimum TE of TDM is longer than that of the SE and ME sequences due to the multiple refocusing pulses applied between the diffusion gradients. The increased in minimum TE (around 17 ms compared to the standard SE or the 1^st^ echo of the ME sequence) will result in approximately a 20% loss of SNR (T2=75 ms). Future work will concentrate on consolidating the three refocusing RFs into one MB3 RF, potentially reducing the TE by approximately 10 ms. Additionally, techniques such as Power Independent of Number of Slices^49^(PINS) can be further explored to mitigate peak power concerns. Despite the implementation of a dedicated slice loop for SE and ME to decrease the TR in the scan protocol, the total scan time remained approximately 2 hours for acquiring the 6-TE data, each comprising 25 diffusion directions, using three sequences. Future work can explore the utilization of spherical b-tensor^50^ encoding to measure and model the distribution of diffusion tensors, potentially yielding further reductions in the overall scan time.

## Conclusion

To conclude, we implemented and systematically compared the multi-echo EPI and TDM sequences for accelerating relaxation-diffusion MRI acquisition. The microstructure analysis of the rdMRI data indicates that TDM can provide a more accurate estimation of T2-relaxation time and microstructure estimation as single-TE EPI, whereas the results from ME are highly biased.

## Data Availability Statement

All the sequences performed in this work and the corresponding reconstruction code are accessible at: https://github.com/QiangLiu0310/Pulseq_TDM_acc_rdMRI. The data that support the findings of this study are available from the corresponding author upon reasonable request.

## Acknowledgments

This study is supported by the National Institutes of Health, Grant Numbers: R01MH116173, R01MH125860, R01NS125781, K01MH117346, R21MH126396, R01EB032378 and VA grant 36C24E22C0027.

